# Discrimination of edge orientation by bumblebees

**DOI:** 10.1101/2022.01.17.476662

**Authors:** Marie Guiraud, Mark Roper, Stephan Wolf, Joseph L. Woodgate, Lars Chittka

## Abstract

Simple feature detectors in the visual system, such as edge-detectors, are likely to underlie even the most complex visual processing, so understanding the limits of these systems is crucial for a fuller understanding of visual processing. We investigated the ability of bumblebees (*Bombus terrestris*) to discriminate between differently angled edges. In a multiple-choice, “meadow-like” scenario, bumblebees successfully discriminated between angled bars with 7° differences, significantly exceeding the previously reported performance of eastern honeybees (*Apis cerana*, limit: 15°). Neither the number of choices required to learn, nor the bees’ final discrimination performance were affected by the angular orientation of the training bars, indicating a uniform performance across the visual field. Previous work has found that, in dual-choice tests, eastern honeybees cannot reliably discriminate between angles with less than 25° difference, suggesting that performance in discrimination tasks is affected by the training regime, and doesn’t simply reflect the perceptual limitations of the visual system. We used high resolution LCD monitors to investigate the limits of bumblebees’ angular resolution in dual-choice experiments. Bumblebee could still discriminate 7° angle differences under such conditions (exceeding the previously reported limit for *Apis mellifera*, of 10°, as well as that of *A. cerana*), eventually reaching similar levels of accuracy, but required longer learning periods than under multiple-choice conditions. Bumblebees show impressive abilities to discriminate between angled edges, performing better than two previously tested species of honeybee. This high performance may, in turn, support complex visual processing in the bumblebee brain.

## Introduction

Low-level feature detectors [1–5] such as edge orientation detector neurons [6] underlie visual object recognition, even in complex cognitive tasks [2,6–10]. Roper *et al.* [11] demonstrated that it is possible to identify and discriminate a wide variety of complex visual patterns, using a low number of edge orientation detectors, without any need for storing “snapshot” visual memories. Differences in edge detection performance are thus likely to underpin interspecies differences in many visual discrimination tasks, so a detailed understanding of visual learning by bees and other insects will require an understanding of the limits of edge orientation detection. Bumblebees are a popular model for studies of insect visual learning [12,13], providing significant advantages in that they can be can be bred and kept in indoor settings, which allows researchers to test them year-round in precisely defined laboratory conditions. There is currently no behavioural data available on how well bumblebees can discriminate between angled edges.

Wehner and Lindauer [14,15] trained European honeybee workers (*Apis mellifera*) to collect food from feeders bearing either vertical, horizontal or 45° black bars. In tests, bees could discriminate the training pattern from one with only a 10° difference in orientation; at 8°, bees’ performance deteriorated; and they failed completely at 5°. Chandra *et al.* [6] trained eastern honeybee workers (*Apis cerana*) to feeders with black stripes of various angular orientations, presented on a white disk. Two tests were used: in one, the bees chose between two cues, one with stripes matching the trained orientation and one in a different orientation; the other test employed a multiple-choice paradigm, in which bees chose between the trained orientation and 11 deviations from it. Honeybees performed better in the multiple-choice situation, successfully choosing the trained orientation over alternatives rotated by 15°, whereas under the dual-choice conditions they only succeeded in differentiating between stripes that differed by 25°. This result gives a clear demonstration of the risks of inferring the limits of discrimination from behavioural performance, since such performance depends not just on what the bee sees, but also on the training procedure. There is mounting evidence that the conditioning procedure plays an important role in how animals perform in visual discrimination tasks [16–19]. Comparative methodological approaches allow us to understand the advantages and limits of each type of protocol [9,20], and provide insight into the ways performance depends not just on sensory limitations but on how information is processed and used in different contexts.

We investigated the ability of bumblebee workers (*Bombus terrestris audax*) to discriminate between cues with differently oriented edges, in both a multiple-choice, meadow-like environment and in a dual-choice set-up, to determine whether their performance differs from that of honeybees and how it is affected by the behavioural context.

## Methods

### Experiments 1 and 2: Multiple-choice (meadow paradigm)

#### Setup and pre-training

We tested bumblebee workers from two commercially bred colonies (Biobest, Belgium). Each colony was kept in a custom-built wooden nest box (280 × 160 × 110 mm high) and connected via a Perspex tunnel to a foraging arena (700 × 700 × 400 mm high) with white painted walls and green laminated paper floor. High frequency fluorescent lighting (TMS 24F lamps with HF-B 236 TLD ballasts, Phillips, Netherland; fitted with Activa daylight fluorescent tubes, Osram, Germany) were used to illuminate the foraging arena; the flicker frequency of the lights was ~42kHz which is well above the flicker fusion frequency for bees [21,22].

During a pre-training phase, bees were allowed free access to the arena where they could forage on *ad libitum* sucrose solution (20% w/w) from ten feeders placed on the arena floor. Each feeder consisted of a sucrose-solution-filled feeding tube (Ø 7mm) placed horizontally at the centre of a vertically aligned, circular disk (Ø 110mm), made of laminated white paper. Each feeder was supported by an attached block weight (15 × 25 × 45 mm high) that allowed the feeder disk to be aligned perpendicular to the floor arena (Figure 1b). Bees foraging from the feeders were individually marked with number tags (Opalithplättchen, Warnholz & Bienenvoigt, Germany) for identification. Numbered bees that were observed to forage in the arena frequently were allowed to advance to the individual training phase. Pollen was provided *ad libitum* into the colony.

**Figure 1.**
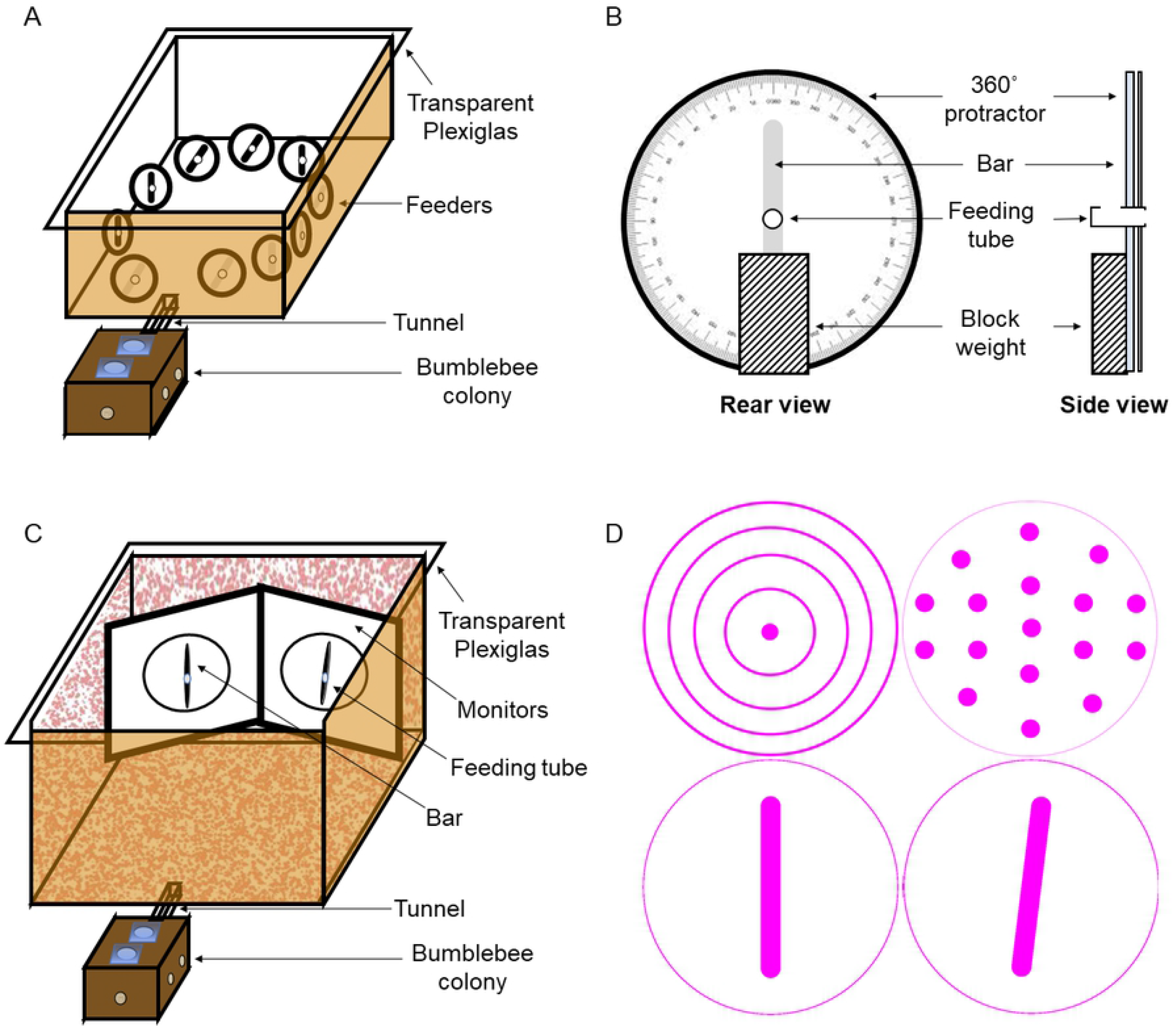
Experimental set-up. **A:** Multiple-choice arena. A bumblebee colony was housed in a wooden box, connected to the experimental arena by a transparent tunnel. Ten feeders were presented in a circle configuration so that all were visible from the centre of the arena. **B:** Feeder. Each feeder consisted of a white paper disc with a printed black bar, and a feeding tube in the centre. A heavy block weight at the rear of the feeder supported a rotatable 360° protractor, allowing the bar to be rotated to the required angle. Five feeders, allocated at random, had the bar oriented to angle *A* and contained a drop of sucrose solution; the others were oriented to angle *B* and contained water only. **C:** Dual-choice arena. Two computer monitors were arranged at a 60° angle at the rear of the arena. Each presented a black bar in a white circle. A feeding tube was placed at the centre of each bar. One bar, allocated at random was oriented to angle *A* and the feeding tube contained a drop of sucrose solution; the other was oriented to angle *B* and contained water. **D:** Example stimuli used in dual-choice experiments. Top row, two stimuli used during pre-training, one on each screen. Bottom row, example bar stimuli used for training and testing: left, 0° bar; right, 7° bar. All stimuli were displayed in magenta (RGB: 255,0, 255) on a white background (RGB: 255, 255, 255).

#### Training and testing

During the training phase, bees were allowed to forage individually in the arena. Ten feeders (Figure 1B) were arranged in a circle, such that a bee could see all ten stimuli from the centre of the arena (Figure 1A). These feeders were identical to those used in pre-training, except that each white circle contained a single black bar (75 x 5 mm). Use of a 360° protractor attached to each feeding station, and spirit level on the arena base, allowed the experimenter to align the bar to precise angular orientations (Figure 1). Five feeders, randomly assigned, had the bar oriented at one angle, *A*, and provided 10μl sucrose solution (CS+: 30% w/w); the remaining five feeders had the bar oriented to a different angle, *B*, and were non-rewarding, filled with a 10μl water droplet (CS-). Feeding tubes were refilled from the rear, preventing sucrose or water from being deposited at the entrance of the tube.

This training paradigm was used for two experiments: in the first, three bees were trained to rewarding and unrewarding feeders with bars differing in orientation by 15°. Each bee experienced one angle *A* (CS+) and one angle *B* (CS-), but three different sets of angles were used, with one set assigned to each bee at random. Angle *A* was either 30°, −60° or −75° (clockwise from vertical); angle *B* was 45°, −75°, or −60° respectively.

In the second experiment, 25 bees were trained to bar orientations differing by 7°. A greater range of bar orientations was used in order to examine discrimination performance across the full range of possible orientations. We started with four base angles at −60°, 0°, 45° and 90°. These were paired with an angle that differed by 7°, either clockwise or anticlockwise (e.g. 45° could be paired with 38° or 52°), giving 8 pairs of angles. We assigned pair of angles to each bee at random and further randomised which of the two angles in the pair was associated with the sucrose reward and which with water. There were thus 16 possible combinations of angles *A* and *B*, although only 12 were assigned to bees in practice. Table 1 gives details of how many bees were trained with each pair of angles.

**Table 1.** Number of bees tested with each pair of bar angles in experiment 2.

| Base angle (°) | Direction of difference | Base angle assignment | Angle A (°) | Angle B (°) | No bees |
| --- | --- | --- | --- | --- | --- |
| -60 | + | A | -60 | -53 | 1 |
| -60 | + | B | -53 | -60 | 2 |
| -60 | - | A | -60 | -67 | 4 |
| -60 | - | B | -67 | -60 | 2 |
| 0 | + | A | 0 | 7 | 1 |
| 0 | + | B | 7 | 0 | 0 |
| 0 | - | A | 0 | -7 | 0 |
| 0 | - | B | -7 | 0 | 0 |
| 45 | + | A | 45 | 52 | 4 |
| 45 | + | B | 52 | 45 | 3 |
| 45 | - | A | 45 | 38 | 2 |
| 45 | - | B | 38 | 45 | 0 |
| 90 | + | A | 90 | 97 | 1 |
| 90 | + | B | 97 | 90 | 1 |
| 90 | - | A | 90 | 83 | 2 |
| 90 | - | B | 83 | 90 | 2 |

Before training, all bees were removed from the arena and the setup was cleaned with 70% ethanol. The focal bee was allowed repeated access to the foraging arena until a total of 100 consecutive feeder choices were recorded. A choice was defined as a bee landing on a feeding tube, and for each choice we recorded whether the feeder was rewarding or unrewarding. After each foraging bout (consisting of multiple feeder choices, typically 4-10, continuing until the bee filled its crop, and returned to the nest box), all feeders were cleaned and the positions of rewarding and unrewarding were randomized before the bee was allowed to re-enter the arena. Rewarding feeders were refilled with 10μl sucrose solution once the bee each time a bee had drunk and departed from the feeder. Bees did not consume water from the unrewarding feeders, so they did not require refilling after each choice, but unrewarding feeders were cleaned and refilled after every foraging bout.

### Experiments 3 and 4: Dual-choice

#### Set-up

We tested bumblebee workers from a third colony (Biobest, Belgium) on dual-choice tests, including even smaller angular deviations. To test the bees’ ability to discriminate below 7°, it is important to ensure that the angular alignment is more precisely controlled than we could guarantee using physical stimuli. Instead, we presented stimuli on two high-resolution, high-refresh-rate LCD computer monitors (Acer Predator GN246HLB, with 144Hz refresh rate, which is above the flicker fusion frequency of bees [22,23]). These monitors were aligned and fixed in position, and software-generated oriented bars were used to ensure uniform angles.

A second flight arena (1150 × 1300 × 500 mm high) was used for experiments 3 and 4. The flight arena was covered with a red Gaussian random dot pattern (generated with custom MATLAB code), printed onto white laser copy paper and laminated. At the top of the arena a fine fabric net was attached, this extended to the laboratory ceiling. High frequency fluorescent lighting (TMS 24F lamps with HF-B 236 TLD ballasts, Phillips, Netherland; fitted with Activa daylight fluorescent tubes, Osram, Germany) were used to illuminate the apparatus.

Two monitor screens were positioned at the rear wall of the flight arena and aligned at 60° angle from each other, allowing the bee to see both screens from the entrance of the arena (980 mm in front of the monitors). A transparent Plexiglas sheet was placed 15 mm from each monitor screen with a small hole (Ø 10mm) at the centre, leading to a feeding tube (Ø 8mm, 15mm long). As the tube was behind the Plexiglas from a bee’s point of view, it did not block any of her flight movements in front of presented stimuli and allowed us an unobstructed view of the bee’s movements.

#### Stimuli

Stimuli were created and displayed on the monitors using custom MATLAB (Mathworks) code and the PsychToolbox [24]. Each monitor displayed a single circle open circle (Ø 260 mm) on a white background, with one or more shapes inside. All stimuli were magenta (RGB: 255,0, 255), allowing observers to easily see the dark body of a bee in front of the monitor while still providing high levels of green-photoreceptor-contrast for the bee, which is required for edge detection and angle discrimination tasks [4,25,26]. During pre-training, one monitor showed a Ø 20 mm filled circle surrounded by four concentric circles (Ø 95, 160, 210 and 260 mm), while the other showed the Ø 260 mm circle containing an arrangement of 17 filled circles (Ø 20 mm; Fig 1D).

During training and tests, each monitor showed a circle containing a single bar (180 mm x 20 mm, with rounded ends; Fig 1D). Each rewarding bar orientation was paired with an identical, unrewarding bar, at a different angle. Two experiments were carried out: in experiment 3, rewarding stimuli were assigned to 6 bees at random (angle *A*: 0°, 38°, 83°, 90°, 90° and 97° clockwise from vertical). The bar for the unrewarding feeder was rotated either +7° or −7°, relative to the rewarding stimulus (angle *B*: 7°, 45°, 90°, 83°, 97° and 90°, respectively). In experiment 4, three bees were trained to stimuli with angle *A* set to 85°, 90° and 90° clockwise from vertical. The bar for the unrewarding feeder was rotated by +5° or −5° relative to the rewarding stimulus (angle *B*: 90°, 85° and 85°, respectively).

#### Pre-training

During the pre-training phase, bees were individually trained to feed from the feeding platforms and to learn that certain visual stimuli were associated with a sucrose reward. During this phase, no oriented bars were presented, but bees learned to discriminate between filled or open circles placed within the 260 mm outer circles (Fig 1D). One stimulus, assigned at random, provided 20μL of 50% (w/w) sucrose solution in the feeding tube (CS+). The other contained 20μL of water (CS-).

A choice was recorded whenever a bee landed on one of the two feeding tubes. If the bee chose the tube in front of the unrewarding stimulus it was allowed to continue making choices it landed on the rewarding feeder. Once the bee had located and consumed the 20μL sucrose solution from the rewarding feeder, it was captured in a transparent ventilated transfer tube (Ø 30mm; 70mm long) and released again from the entrance of the arena to make another choice. Each foraging bout consisted of approximately four feeder visits, and lasted until the bee’s crop was full and it returned to the nest (mean crop capacity: 80 to 120μL, [27]). The positions of the rewarding and unrewarding stimuli were pseudo-randomized between bouts (with no more than two consecutive visits permitted with the same positions). Once a bee chose the rewarding pattern eight times out of the last ten feeder visits, it progressed to the experimental training phase.

#### Training and testing

The experimental training phase followed the same procedure as above. During each foraging bout, each of the two monitors showed an oriented bar (one feeder providing reward of 20μL of 50% sucrose solution w/w, and the other providing 20μL water). The location of the rewarding stimulus was pseudo-randomized for each foraging bout. Each bee was trained for approximately 150 choices, continuing until the bee achieved at least 80% correct choices over two consecutive 10-choice batches. If the bee did not achieve this 80% accuracy after 250 choices, the training was abandoned.

If the bee achieved ≥80% correct choices in two consecutive batches of 10 choices, it was subjected to a test bout. During tests, both feeding tubes had a 20μL drop of water within them regardless of the stimulus. After each batch of 10 choices between unrewarding stimuli, the bee was given 10 choices under training conditions, with the trained stimulus again rewarded with sucrose, to maintain its motivation to visit the feeders. The proportion of choices for the trained stimulus was recorded for each batch of 10 choices for unrewarded stimuli. The test bout ended when the bee no longer attempted to visit either feeding tube.

#### Analysis

Learning performance was evaluated using the percentage of correct choices for every block of 10 consecutive feeder visits. Individual bees were categorised as having learned the task if they made more than 80% correct choices over two consecutive 10-choice blocks at any point during the learning phase. Experiments 1 and 4 had small sample sizes (of only 3 bees per experiment), so we evaluated bees’ performance based solely on how many had reached this learning criterion.

For experiments 2 and 3, we further tested whether the number of correct choices over the last 20 feeder visits by each bee, during training, was significantly greater than chance performance, using one-sample Wilcoxon signed-rank tests to compare the median number of correct choices to a chance level of 10 correct choices out of 20. The bees in experiment 3 were also tested on their performance in unrewarding probe s. We tested whether their performance exceeded chance in these tests using a paired Wilcoxon signed-rank test, in which the number of correct choices made was compared to an expected value of half the total number of feeder visits during the test.

We investigated whether there was any effect of bar orientation on the final performance reached by each bee in experiment 2. Bees were assigned to one of four orientation groups, in which we tested their ability to differentiate between bars at one of four angles (−60°, 0°, 45°, 90°) and bars ±7° from each of those angles (Table 1). Only one bee was tested in the 0°±7° group, so it was excluded from this analysis. The other three groups (−60°, 45°, 90°) contained 9, 9 and 6 bees, respectively. We tested for differences in performance between these groups using a Kruskal-Wallis test, in which the dependent variable was the number of correct choices over the last 20 feeder visits made by bees in each orientation group. Among those bees that reached a performance of 80% correct choices over at least 20 visits, we tested whether there was an effect of bar orientation on the speed of learning, using another Kruskal-Wallis test in which the dependent variable was the number of choices required to reach the 80% learning criterion by bees in each orientation group.

We investigated whether there were differences in final performance between bees trained to discriminate 7° orientation differences under dual- and multiple-choice conditions, using a pair of Kruskal-Wallis tests to compare the performance of bees from experiment 2 to those from experiment 3. The dependent variables were the number of correct choices by each bee during feeder visits 80-100 and the number of correct choices by each bee over its last 20 choices. We tested whether there was a difference in the speed of learning between experiments 2 and 3, using another Kruskal-Wallis test in which the dependent variable was the number of choices required to reach the 80% learning criterion by bees in each experiment.

All statistical tests were 2-tailed. All statistics were calculated in Matlab (version R2018a; Mathworks Inc., Natick, USA).

## Results

### Experiment 1: Multiple-choice, 15° angular difference

The first experiment tested bumblebees’ ability to discriminate a 15° difference in edge orientation. The first three bees tested all reached the criterion for success (mean ≥80% correct choices over two consecutive blocks of 10 feeder visits) within 40-50 feeder choices (proportion correct choices over final 20 choices, mean ± standard error of the mean: 0.917 ± 0.033; Fig 2). Due to this clear evidence that bees could discriminate an angular difference of 15°, we proceeded immediately to test a smaller difference of 7°.

**Figure 2.**
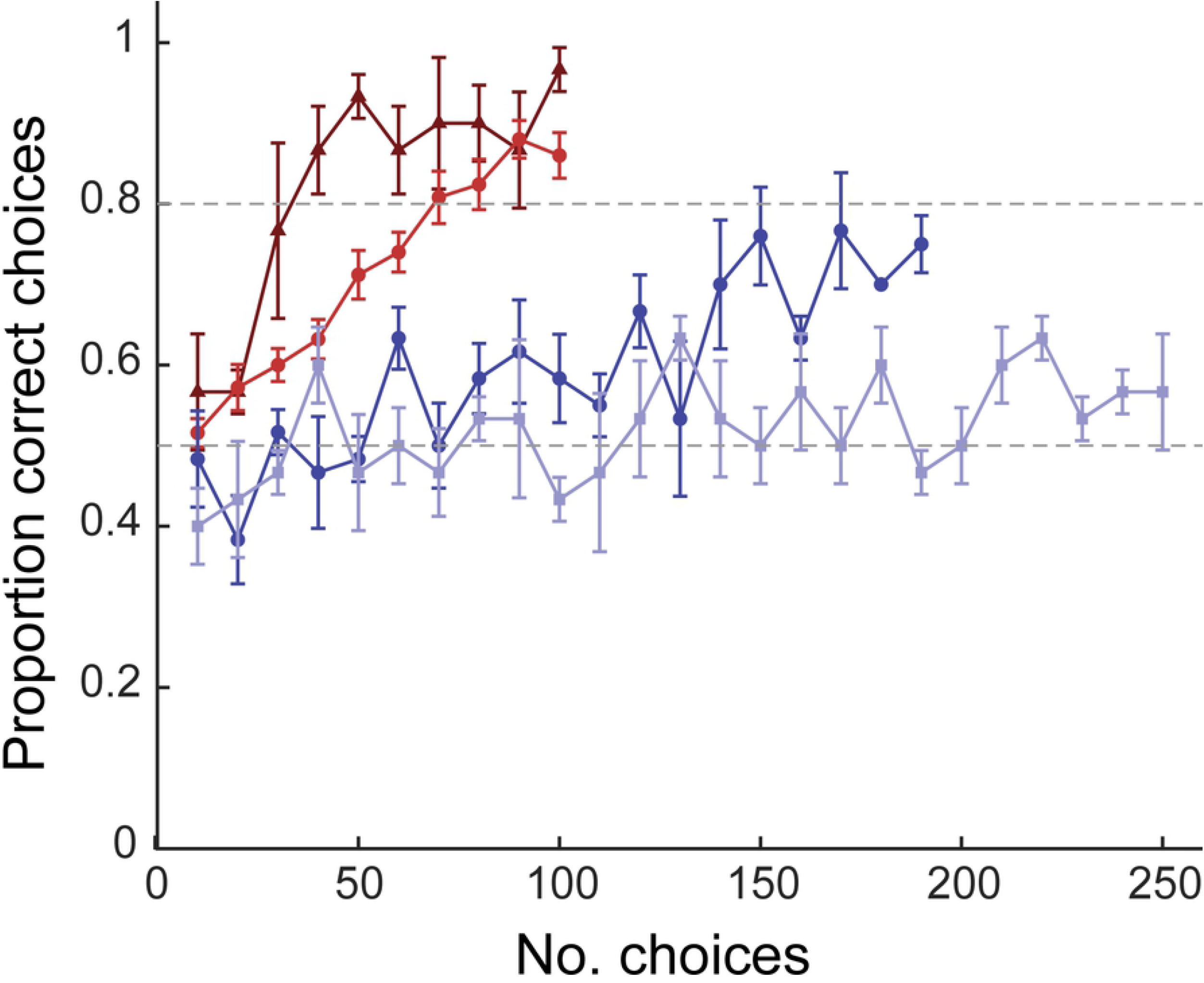
Learning curves for bees in multiple- and dual-choice experiments. Each marker shows the mean proportion of choices (±S.E.) for the trained angle, *A*, across a block of 10 consecutive choices in multiple-choice experiments (red lines) and dual-choice experiments (blue lines). Dark red triangles indicate experiment 1 (multiple-choice, angles *A* and *B* differ by 15°, N=3); light red circles, experiment 2 (multiple-choice, angles *A* and *B* differ by 7°, N=25); dark blue circles, experiment 3 (dual-choice, angles *A* and *B* differ by 7°, N=6); light blue squares, experiment 4 (dual-choice, angles *A* and *B* differ by 5°, N=3). Dashed grey horizontal lines indicate a chance level of performance (0.5) and the criterion for our bees to be classified as having learned (0.8, but note that bees had to reach this proportion of correct choices over 20 feeder visits).

### Experiment 2: Multiple-choice, 7° angular difference

Out of 25 bees tested on angled bars with a 7° difference, just three failed to reach 80% correct choices over at least two consecutive blocks of 10 feeder visits. Those bees that reached the 80% learning criterion, did so after a mean of 71.36 ± 3.37 choices (range: 40-100; Fig 2). The mean proportion of correct choices by all 25 bees in this group over their last 20 feeder visits was 0.870 ± 0.025, significantly greater than expected by chance (Wilcoxon signed-rank test: V = 325, N = 25, P <0.0001), and similar to the performance seen in the 15° experiments (Fig 2).

The three bees that failed to reach 80% successful choices were each trained with a different bar orientation, *A* (−60°, 52° and 45°) and each of these angles was also assigned to other bees that did reach the criterion (5, 2 and 5 successful learners, respectively). There were no significant differences between the groups of bees trained on each of three different base bar orientations (−60°±7°, 45°±7°, 90°±7°) in the number of correct choices over their final 20 choices (Kruskal-Wallis test: χ^2^_2_ = 4.98, P = 0.083; Fig 3A-B). Among the 21 bees in these three orientation groups that did reach the 80% learning criterion, there was no effect of bar orientation on the number of choices required to reach this level of performance (Kruskal-Wallis test: χ^2^_2_ = 1.60, P = 0.450; Fig 3C).

**Figure 3.**
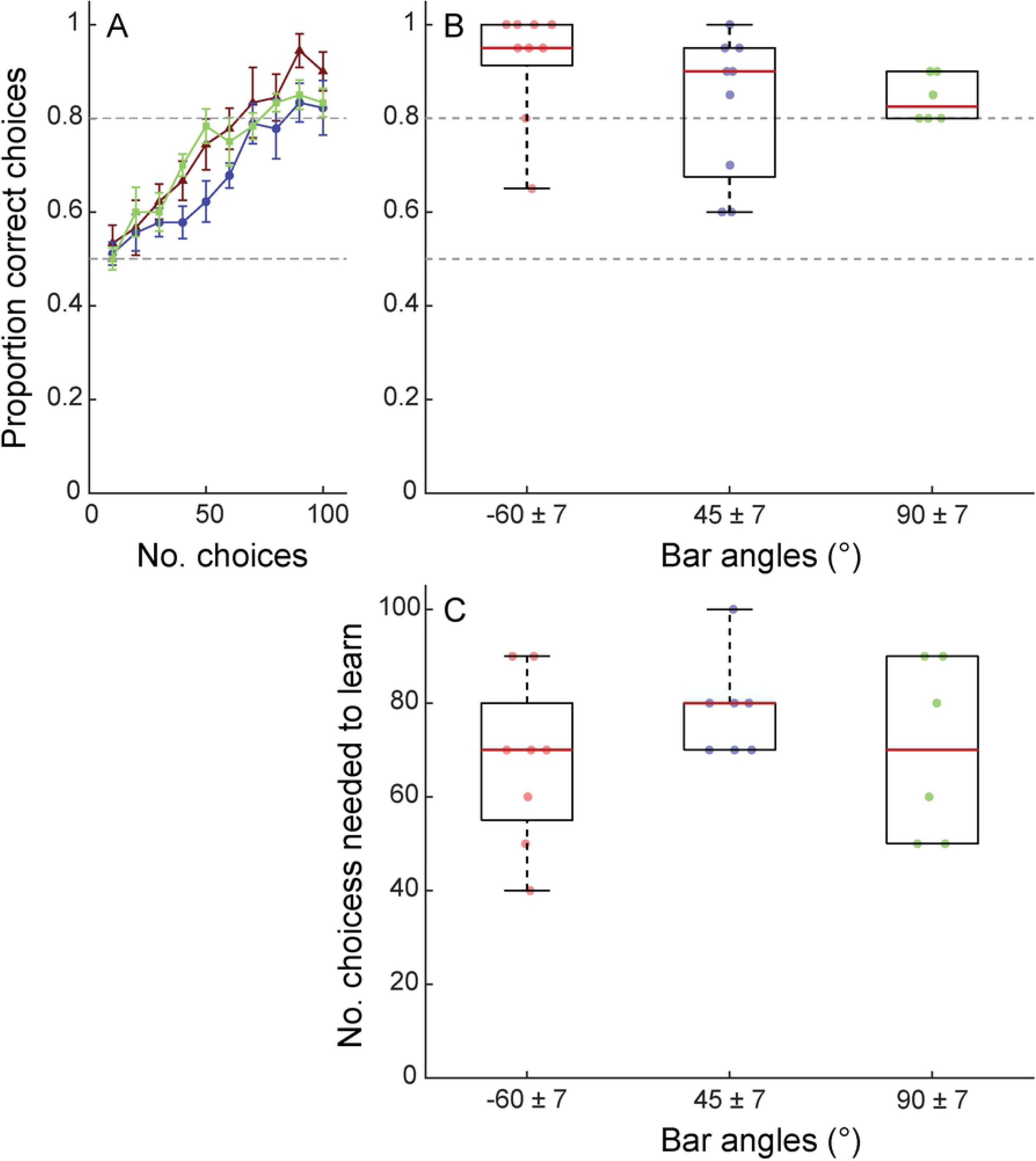
Angular discrimination is unaffected by training bar angles, in multiple-choice tests. **A:** Mean proportion of choices (±S.E.) for the trained angle, *A*, across a block of 10 consecutive choices for bees in experiment 2. Dark red triangles indicate bees that were trained to discriminate bars of −60° ± 7° (N=9); blue circles, bees trained to discriminate bars of 45° ± 7° (N=9); light green squares, bees trained discriminate bars of 90° ± 7° (N=6). Dashed grey horizontal lines indicate a chance level of performance (0.5) and the criterion for our bees to be classified as having learned (0.8 but note that bees had to reach this proportion of correct choices over 20 feeder visits). **B:** Proportion of choices for angle *A* (correct choices) across the last 20 feeder visits by bees trained to each group of angular orientations (−60° ± 7°, 45° ± 7°, 90° ± 7°). Red lines indicate group median, boxes indicate the interquartile range and whiskers indicate the range. Filled circles show the proportion of choices for angle *A*, made by each individual. There are no significant differences between groups. **C:** Number of feeder visits required to reach a performance level of at least 80% correct choices across a block of 20 consecutive feeder visits, by bees trained to each group of angular orientations (−60° ± 7°, N=8; 45° ± 7°, N=7; 90° ± 7°, N=6). Three bees that never reached this criterion have been excluded (one from the −60° ± 7° group and two from the 45° ± 7° group). There are no significant differences between groups.

### Experiment 3: Dual-choice, 7° angular difference

We also tested bees’ ability to discriminate a 7° angle variance in a dual-choice paradigm. After 100 feeder visits (the total number experienced by bees in the multiple-choice experiments), six bees had a mean success rate of just 0.600 ± 0.058 across choices 80-100. This performance is significantly lower than in the multiple-choice setup (Kruskal-Wallis test: χ^2^_1_ = 10.41, P = 0.0013; Fig 2).

With continued training, all but two bees reached the 80% criterion within 130-170 choices (mean ± S.E.: 147.50 ± 8.54), demonstrating that bumblebees are capable of learning to discriminate angular deviations of 7° under these conditions. Bees in the dual-choice experiment required significantly more training to reach this criterion than those in the multiple-choice experiment (Kruskal-Wallis test: χ^2^_1_ = 9.99, P = 0.0016), and examination of the learning curves makes it clear that bees’ performance in experiment 3 improved at a lower rate than that of bees in experiment 2, throughout the entire training period (Fig 2).

Even the two bees that never reached the 80% threshold showed evidence of learning, with final performances of 0.70 and 0.75 after 190 choices. The mean proportion of correct choices by all 6 bees in this group over their last 20 feeder visits was 0.800 ± 0.034, significantly greater than expected by chance (Wilcoxon signed-rank test: V = 21, N = 6, P = 0.031; Fig 2). There was no significant difference between the final performance reached by these bees and those discriminating 7° differences in experiment 2, under multiple-choice conditions (Kruskal-Wallis test: χ^2^_1_ = 2.72, P = 0.099).

The bees in this test were subjected to a learning test after they had reached the 80% criterion, or after 190 choices in the case of the two bees that never reached 80% performance. In this test, all feeders were unrewarding. Under these conditions, bees were as successful in discriminating angles as they were at the end of the training period, with a mean proportion of choices for the previously rewarding stimulus of 0.764 ± 0.065, significantly greater than expected by chance (Wilcoxon signed-rank test: V = 21, N = 6, P = 0.031).

### Experiment 4: Dual-choice, 5° angular difference

We attempted to train a further three bees to discriminate oriented bars with only 5° angle differences. This sample size is too low for formal analysis, but Fig 2 suggests that these bees did show some improvement in performance with continued training. However, they appeared to improve in performance at a far slower rate than those discriminating 7° differences. Even after 250 feeder visits over the course of four to six full days, the bees did not obviously choose the correct bars more often than expected by chance (mean proportion correct choices ± S.E.: 0.600 ± 0.029; individual means: 0.500, 0.600, 0.650; Fig 2). Performance in a subsequent learning test was similarly poor (mean proportion correct choices ± S.E.: 0.636 ± 0.016; individual means: 0.625, 0.667, 0.615).

## Discussion

In this study, we demonstrate that bumblebees (*Bombus terrestris*) are able to discriminate between oriented bars with an angle difference of just 7°, in both a meadow-like multiple-choice, and a dual-choice scenario, and regardless of bar orientation. Our data suggest that bumblebees are unlikely to be able to discriminate angular differences of 5° or less. This performance slightly exceeds that reported for European honeybees (*Apis mellifera*), which discriminated 10° differences with certainty and still showed some evidence of discrimination at 8°, in dual choice tests [14,15]. By contrast, previous work has found that eastern honeybees (*Apis cerana*) could not be trained to discriminate angle differences below 15° in a multiple-choice setup, and below 30° under dual-choice conditions [6].

While the final performance of bumblebees was similar in both experiments, it took far more training for them to reach a high level of performance when choosing between two options displayed on computer monitors than when 10 options were displayed in a physical array. A number of studies have demonstrated an influence of conditioning paradigm on bees’ behavioural performance in a variety of tasks [28–30]. Our study, in concert with that of Chandra *et al.* [6], demonstrates that the training procedure can affect the outcome of even apparently simple perceptual tasks and highlights the dangers of attempting to derive perceptual limits directly from behavioural studies.

Why might bees learn to discriminate angles more quickly in a multiple-choice experiment? There were several methodological differences between the experiments, so it is not possible to determine this conclusively, but one important factor may be the frequency with which bees were able to get feedback on their choices. In the meadow-like, multiple-choice experiment, bees visited several feeders during every foraging bout (round-trip from the nest to the flight arena and back), but in the dual-choice experiment they could make only one choice before being returned to the arena entrance. The comparative difference in how rapidly they can sample different options and rewards may account for the difference in learning speed.

The stimuli for our multiple-choice test were printed on paper, while the dual-choice test was presented on computer monitors, which have previously been found make fine discriminations more difficult for bees [31]. However, Chandra et al. [6] reported differences between dual- and multiple-choice setups for eastern honeybees, even though their stimuli were printed on paper in both experiments, so the use of monitors is unlikely to account for the differences we observe. Chandra et al. [6] suggested that the variation in performance was due to their multiple-choice configuration providing only a one in twelve chance of the bee selecting the correct stimulus. Under dual-choice conditions it may be efficient to sample all stimuli at random, rather than investing time and computation in discriminating between them, since there is a 0.5 chance of reward from each feeder sampled. By contrast, when the odds of success fall to 0.0833, an investment in learning to discriminate between correct and incorrect stimuli may lead to a greater medium- or long-term rate of energy gain. The odds of success from random sampling were 0.5 in both of our experiments, however, so differences in expected reward from random sampling cannot account for the difference between paradigms.

While our results do show that aspects of the training procedure can affect bees’ performance in angular discrimination tasks, we can nonetheless draw some conclusions regarding the sensory/perceptual limitations faced by bumblebees. At the end of training, when bees had reached saturation-level performance, both groups of bees discriminated between edges differing by 7° with a high level of accuracy, clearly demonstrating that they could perceive differences of at least that magnitude. When trained to bars with a 5° difference under the dual-choice paradigm, three bees did not perform above chance levels even after an extra 2-3 days of training and 100 more choices than were required to learn a 7° difference. A greater sample size would be required to confirm that bees are unable to learn a 5° difference, but given the slow rate of improvement and the fact that bees typically invest relatively little time in learning how to handle flowers [32–34], we suggest that the practical limit for bumblebee angular perception during foraging tasks is likely to lie between 5°-7°. Note that it is difficult to derive conclusions about perceptual limits from behavioural data: discriminating 5° differences may not actually be beyond bees’ perceptual ability, just sufficiently difficult or costly that doing so was less efficient than random sampling in our experiment.

It is interesting to note that, even in a multiple-choice setup, the limit of angular discrimination previously reported for eastern honeybees [6] is double what we found for bumblebees, pointing to a significant variation in visual discrimination ability between the two species. One explanation might be that visual-spatial resolution is constrained by eye optics: bumblebee workers are generally larger than honeybees and their larger eyes have both larger ommatidial facets and reduced interommatidial angles [35–39]. As a result, bumblebees have been found to have higher visual acuity than European honeybees (*Apis mellifera*), in several behavioural contexts [23,36,37]. Recent work on X-ray micro computed-tomography of the honeybee and bumblebee eyes [37,38,40,41] may pave the way toward future experiments in which we can relate the behavioural performance of individual insects directly to the acuity of their eyes. Bumblebee workers exhibit a high degree of variation in body size, so a fruitful avenue of investigation may be to ask whether larger individuals, with larger eyes, perform better in angular discrimination tasks. In these experiments we did not take any measures of body size, but we took care to use individuals that were within the normal size range for bumblebee workers and the bees that failed to reach a high level of performance in our experiments did not appear notably smaller than successful bees.

European honeybees have been reported to show levels of angular discrimination only a little less than those we found in bumblebees [15]. The size and morphology of *A. mellifera* eyes lies between that of *A. cerana* and *B. terrestris* in many aspects (e.g. ommatidial number: *A. cerana* = ~4921, *A. mellifera* = ~5375, *B. terrestris* = ~5656; eye surface (mm2): *A. cerana* = 2.3, *A. mellifera* = 2.5, *B. terrestris* = 2.8; [39,42]), yet European honeybees performance was closer to that of bumblebees than this might predict. This perhaps suggests that optical resolution is not the limiting factor on honeybee performance; the characteristics of the neurons mediating edge orientation discrimination are also likely to play an important role.

Our results and those of Chandra et al. [6], show that bees could discriminate angular differences regardless of bar orientation. Chandra et al. [6] used a mathematical model to suggest that a minimum of three orientation sensitive neurons would be required to account for this orientation indifference. However, this model assumed neurons with horizontal and vertical maximal sensitivities. Empirical work has subsequently identified two types of edge orientation sensitive neurons in the lobula, the third visual ganglion of the honeybee, with maximal sensitivity to edges angled at 115° and 220° from the vertical, respectively [43]. Roper *et al.* [11] presented computational-neuronal models based on the known properties of honeybee [43] and dragonfly neurons [44], which predicted performances remarkably similar to empirical honeybee results. Both models predicted that discrimination performance should be independent of the orientation of the training bars, as found in all three species of bees so far tested. These models support the hypothesis that angular discrimination performance is largely determined by the very limited number of orientation detector neurons identified in insects, and that these neurons’ angular tuning is adaptive in allowing uniform performance across a wide variety of orientations.

## Acknowledgments and funding

This study was supported by Human Frontier Science Program grant (RGP0022/2014; www.hfsp.org) to L.C.; Engineering and Physical Sciences Research Council program grant Brains-on-Board (EP/P006094/1; epsrc.ukri.org) to L.C.; European Research Council grant SpaceRadarPollinator (339347; erc.europa.eu) to L.C.; and a Royal Society Wolfson Research Merit Award (royalsociety.org) to L.C. The funders had no role in study design, data collection and analysis, decision to publish, or preparation of the manuscript.

## Authors’ contributions

Conceptualization, M.R., M.G., S.W. and L.C.

Methodology, M.R., M.G. and S.W.

Investigation, M.R., M.G. and S.W.

Data Curation, M.G.

Software, M.G.

Formal analysis, M.G. and J.L.W.

Visualization, M.G. and J.L.W.

Writing - Original draft, M.G.

Writing – Review & editing, M.G., M.R., J.L.W. and L.C.

Funding acquisition, L.C.

Supervision, L.C.

